# Psychometric properties of the PHQ-9 depression scale in people with multiple sclerosis: a systematic review

**DOI:** 10.1101/321653

**Authors:** Sarah Patrick, Peter Connick

## Abstract

**Background:** Depression affects approximately 25% of people with MS (pwMS) at any given time. It is however under recognised in clinical practice, in part due to a lack of uptake for brief assessment tools and uncertainty about their psychometric properties. The 9-item Patient Health Questionnaire (PHQ-9) is an attractive candidate for this role.

**Objective:** To synthesise published findings on the psychometric properties of the 9-item Patient Health Questionnaire (PHQ-9) when applied to people with multiple sclerosis (pwMS).

**Data sources:** PubMed, Medline and ISI Web of Science databases, supplemented by hand-searching of references from all eligible sources.

**Study eligibility criteria:** Primary literature written in English and published following peer-review with a primary aim to evaluate the performance of the PHQ-9 in pwMS.

**Outcome measures:** Psychometric performance with respect to appropriateness, reliability, validity, responsiveness, precision, interpretability, acceptability, and feasibility.

**Results:** Seven relevant studies were identified, these were of high quality and included 5080 participants from all MS disease-course groups. Strong evidence was found supporting the validity of the PHQ-9 as a unidimensional measure of depression. Used as a screening tool for major depressive disorder (MDD) with a cut-point of 11, sensitivity was 95% sensitivity and specificity 88.3% (PPV 51.4%, NPV 48.6%). Alternative scoring systems that may address the issue of overlap between somatic features of depression and features of MS *per se* are being developed, although their utility remains unclear. However data on reliability was limited, and no specific evidence was available on test-retest reliability, responsiveness, acceptability, or feasibility.

**Conclusions:** The PHQ-9 represents a suitable tool to screen for MDD in pwMS. However use as a diagnostic tool cannot currently be recommended, and the potential value for monitoring depressive symptoms cannot be established without further evidence on test-retest reliability, responsiveness, acceptability, and feasibility.

**PROSPERO register ID:** CRD42017067814

## Introduction

Multiple sclerosis (MS) is estimated to affect over 2.3 M people globally(1). It is a chronic inflammatory and degenerative disease of the central nervous system that typically results in sensory, motor, and cognitive impairments(2). The potential co-existence of depression in people with MS (pwMS) is well recognised, with a lifetime prevalence >50%, point prevalence of approximately 25%(3), and doubling of the standardized mortality rate for suicide compared to the general population(4). Depression in pwMS is nevertheless underdiagnosed in clinical practice(5) despite being a major determinant of quality of life(6), and responsive to standard therapeutic approaches(7). This in part may reflect uncertainty around the optimum approach to evaluation. A ‘gold-standard’ approach based on the Structured Clinical Interview for Diagnostic and Statistical Manual of Mental Disorders is impractical at the necessary scale of clinical practice. Self-reporting through use of patient reported outcome measures (PROMS) therefore provides an attractive option for screening, and as a potential approach for monitoring in both clinical and research settings.

A number of PROMS have been applied to quantify the burden of affective symptoms in pwMS across research and clinical settings, including the Beck Depression Inventory (BDI-II), Hospital Anxiety and Depression Scale (HADS), and the nine-item Patient Health Questionnaire (PHQ-9)(8). The PHQ-9 scale is notable in this context because it is freely available and has been validated across a wide range of clinical populations(9). The PHQ-9 is a self-report version of the Primary Care Evaluation of Mental Disorders (PRIME-MD), developed in the mid-1990s by Pfizer Inc.(10) It evaluates depressive symptoms over the preceding two weeks, with 9-items each allowing four response-options for the frequency of symptom-experience. A total score is derived by summation, and interpreted against established thresholds. Despite the potential advantages of the PHQ-9 as a tool to evaluate depressive symptoms in pwMS, there have been no previous focused reviews of the PHQ-9’s psychometric performance in this population.

Fitzpatrick *et al.* propose a framework for the evaluation of patient reported outcome measures (PROMs) such as the PHQ-9 based on eight key performance indicators(11). These are: *Appropriateness* for the specific role intended, such as screening, diagnosis, or monitoring; *Reliability* ; *Validity; Responsiveness* –whether an instrument is sensitive to changes of importance to patients; *Precision* – the number and accuracy of distinctions made by an instrument; *Interpretability* of scores; *Acceptability* to respondents of using the instrument; and *Feasibility* for deployment in clinical practice or research. The aim of this review was to evaluate the known performance of the PHQ-9 against these performance criteria.

## Methods

Design of the systematic review was based upon PRISMA guidelines and we used the PRISMA checklist when writing our report (12). The study protocol was documented in advance on the PROSPERO database (https://www.crd.york.ac.uk/PROSPERO;/ S1 fig.). We used the PRISMA checklist when writing our report.

## Information Sources & search strategy

Evidence was gathered from the databases ‘PubMed’, ‘Medline’ and ‘ISI Web of Science’, supplemented by hand-searching of references from all eligible sources. Search terms used were ‘Multiple Sclerosis’ ‘PHQ-9’, and the related terms (‘MS’, ‘Disseminated Sclerosis’, ‘PHQ Patient Health Questionnaire’, ‘Patient Health Questionnaire 9’, ‘PRIME-MD’).

## Eligibility Criteria and study selection

After gathering the evidence, the following eligibility criteria were applied. The sources were required to be primary literature written in English and published following peer-review with a primary aim to evaluate the performance of the PHQ-9 in pwMS. Studies that simultaneously evaluated other depression inventories or other conditions were considered to be eligible. No date restriction on eligibility was applied. Initial screening of abstracts was performed by a single author (SP). Full articles were then retrieved and eligibility assessment performed, with a final decision over study inclusion taken in consensus with a second reviewer (PC).

## Data Collection

Data were extracted by a single author (SP) using a standardised form that captured details about the study (authors, year, country), the samples (size, diagnoses, method of recruitment, baseline demographic characteristics), and ‘quality indicators’ as defined by the STrengthening the Reporting of OBservational studies in Epidemiology (STROBE) checklist (S2 Fig.)(13). Summary measures were also extracted for the eight performance indicators as described below.

## Risk of bias in individual studies

As no statistical synthesis was planned, quality assessment was conducted for the purposes of describing the conduct of the included studies. SP assessed the included studies for methodological quality based on criteria defined by the STROBE checklist.

## Summary Measures

Relevant measurement with regards to the eight performance indicators was pre-defined as follows. Appropriateness was defined by identification of whether the PHQ-9 was being tested as a screening, diagnostic, or monitoring tool for depression/suicidality in pwMS. Reliability was defined by evaluation of internal (e.g. split-half / Cronbach’s alpha) and external measures (test-retest). Validity was defined based on criterion, concurrent and discriminant approaches. Studies that attempted to define the dimensionality of the PHQ-9 were also interpreted to represent validity studies for the underlying constructs of depression and suicidality. Responsiveness was defined as a determination of QOL change and/or therapeutic response. Precision was defined as exploration of alternative scoring paradigms and evaluation of their relative utility. Interpretability of scores was defined as using ecological validation approaches and/or relationships to QOL or other depression indicators. Acceptability was defined as the collection of participant feedback either quantitative or qualitative. Feasibility was broadly interpreted as data on practical aspects of administration such as completion rates, time to complete, suitability for various subpopulations (e.g. sensory impaired etc.).

## Synthesis of results

A narrative synthesis was used to describe findings on the eight psychometric performance indicators.

## Risk of bias across studies

Selective reporting was evaluated by the ‘STROBE’ quality assessment. Potential overlap of cohorts between publications was identified by consideration of authorship and cohort characteristics.

## Results

## Study Selection

One hundred and sixteen titles were identified by initial search. After elimination of duplicates, 58 unique citations remained, of which 49 were excluded by screening the title. Nine articles were included for assessment of full-text, of which one was excluded due to publication only in abstract form, and a further study was excluded due to being a secondary analysis of previously published data. Seven studies were therefore included in the review (Fig 1).

**Fig 1.**
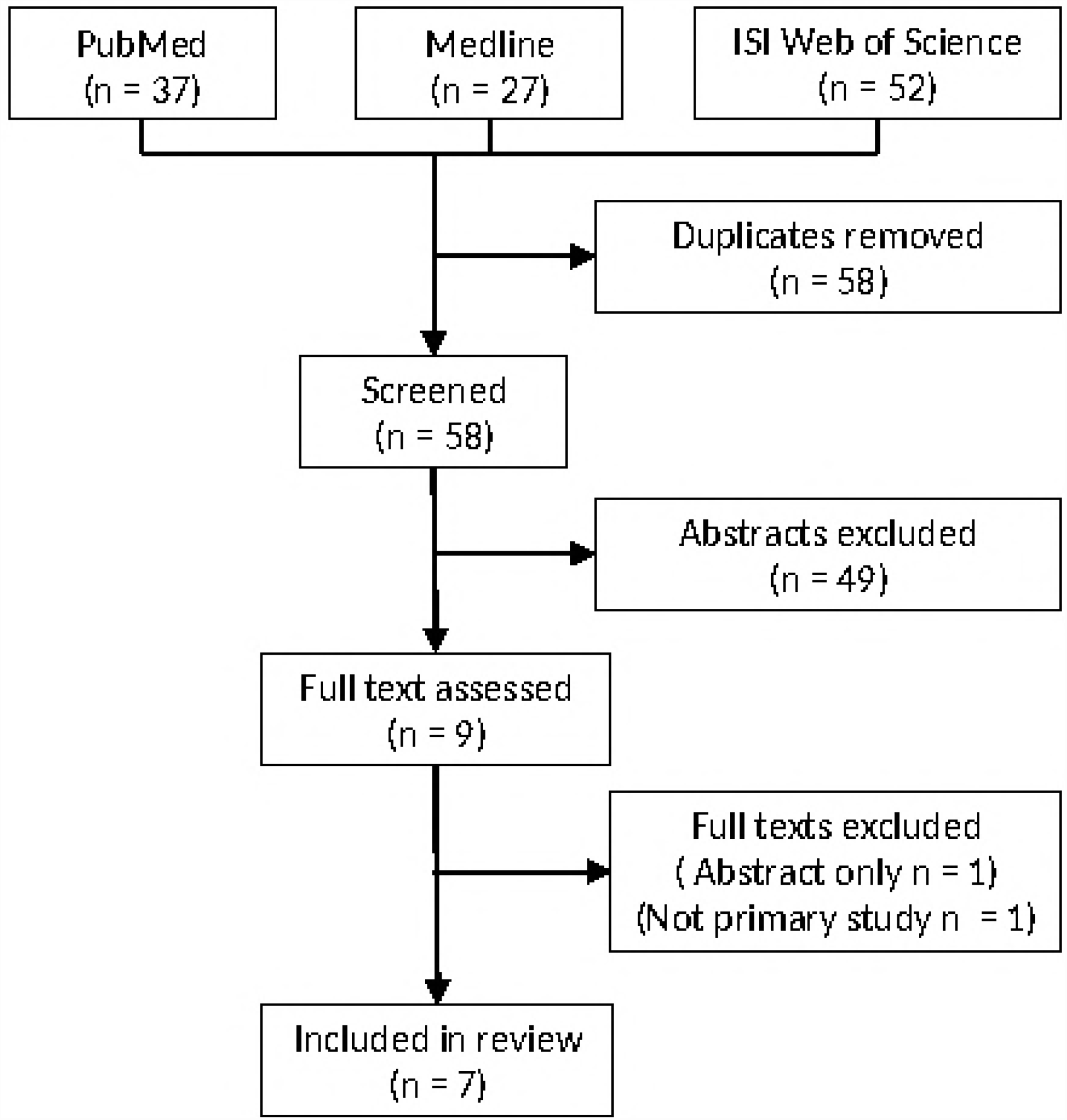
Study selection flow diagram.

## Study Characteristics

Included studies were published between 2012 and 2017 by research groups based in the USA (n=4) (14–17) and Canada(n=3) (18–20). Overall study quality was high (mean quality score 75%, range 62.5 to 93.8; Table 1). Two quality items were consistently low scoring across all studies; prospective definition and/or mitigation of potential bias, and prior sample size calculations. A total of 5080 individuals were included in our review, although highly similar cohort characteristics between Amtmann *et al.* (2014)(14) and Amtmann *et al.* (2015)(15), also between Patten *et al.(19) and Altura et al.(20) were apparent and* raised the possibility that a number of participants (up to 12%) had been analysed twice (Table 2). Separately, one large study (n=3507; Gunzler *et al.* (16)) provided 69% of the total number of participants. Excluding this study, the potential rate of double analysis rose to 39%.

**Table 1.** STROBE checklist quality metrics for included studies.

|  | Sjonnese<br>(2012) | Amtmann<br>(2014) | Amtmann<br>(2015) | Patten<br>(2015) | Gunzler<br>(2015) | Altura<br>(2016) | Kim<br>(2017) |
| --- | --- | --- | --- | --- | --- | --- | --- |
| Objective(s) described | 1 | 1 | 1 | 1 | 1 | 1 | 1 |
| Design described | 0.5 | 0.5 | 1 | 0.5 | 0.5 | 0 | 0.5 |
| Setting described | 1 | 1 | 1 | 1 | 1 | 1 | 0.5 |
| Participant selection described | 0.5 | 0.5 | 1 | 1 | 0.5 | 1 | 0.5 |
| Recruitment methods described | 1 | 1 | 1 | 1 | 0 | 1 | 0 |
| Assessment schedule described | 1 | 1 | 1 | 1 | 1 | 1 | 1 |
| Key cohort characteristics included | 0.5 | 0.5 | 0.5 | 0.5 | 0.5 | 0.5 | 0.5 |
| Data capture methods described | 0.5 | 1 | 1 | 1 | 0.5 | 1 | 0 |
| Bias addressed | 0 | 0 | 0 | 1 | 1 | 1 | 0 |
| Prior sample size calculation | 0 | 0 | 0 | 1 | 0 | 0 | 0 |
| Prior data handling plan | 1 | 1 | 1 | 1 | 1 | 1 | 1 |
| Statistical analysis plan | 1 | 1 | 1 | 1 | 1 | 1 | 1 |
| Description of recruited and non-recruited subjects | 1 | 1 | 1 | 1 | 0 | 1 | 1 |
| Description of cohort characteristics | 0.5 | 1 | 0.5 | 1 | 1 | 0.5 | 1 |
| Report of data completeness | 0.5 | 1 | 1 | 1 | 0 | 0.5 | 1 |
| Main results provided | 1 | 1 | 1 | 1 | 1 | 1 | 1 |
| **Total Score** | 11 | 12.5 | 13 | 15 | 10 | 12.5 | 10 |

**Table 2.**
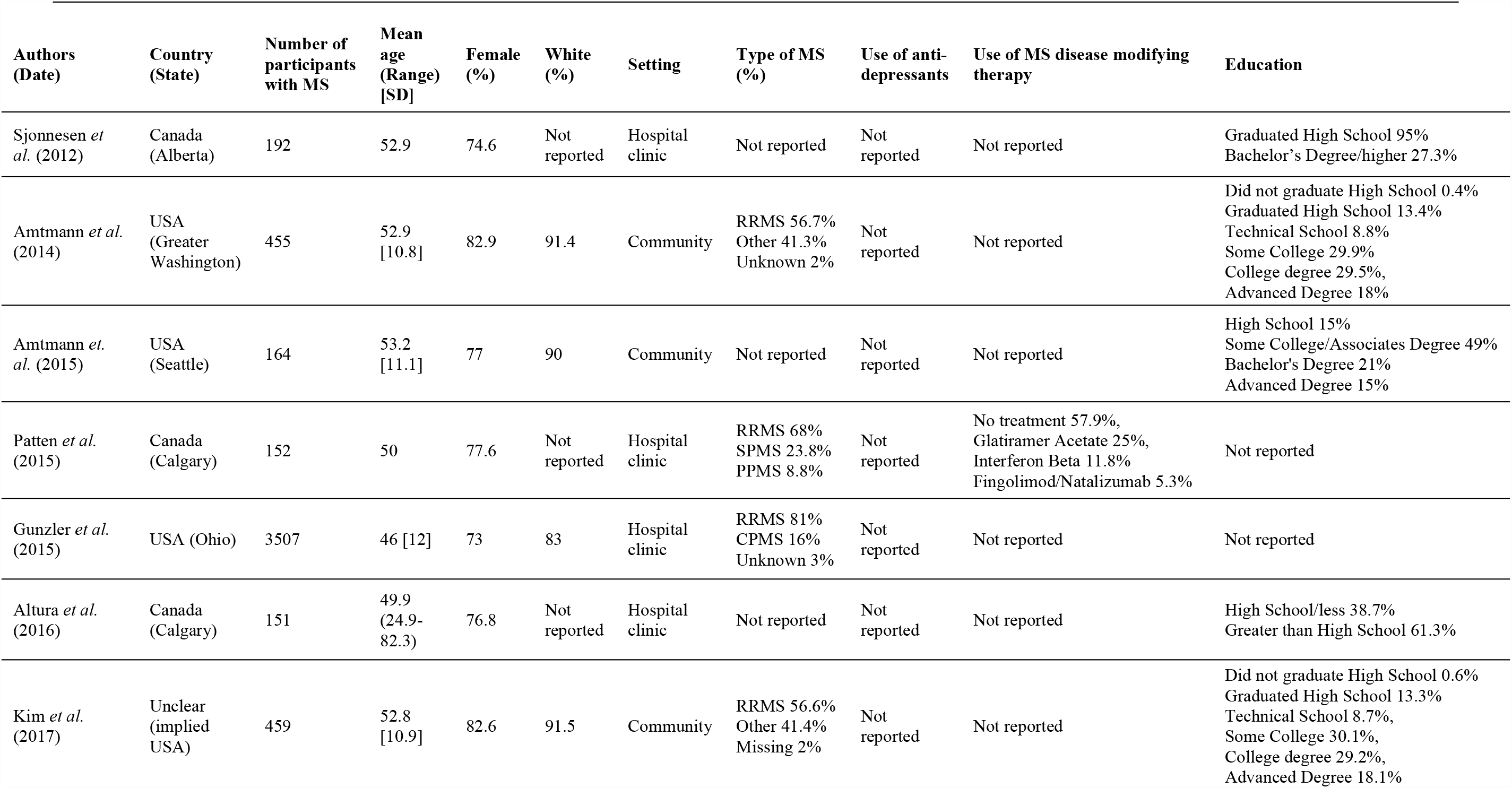
Cohort characteristics for included studies.

| Authors<br>(Date) | Country<br>(State) | Number of<br>participants<br>with MS | Mean<br>age<br>(Range)<br>[SD] | Female<br>(%) | White<br>(%) | Setting | Type of MS<br>(%) | Use of anti-<br>depressants | Use of MS disease modifying<br>therapy | Education |
| --- | --- | --- | --- | --- | --- | --- | --- | --- | --- | --- |
| Sjonnese *et al.* (2012) | Canada<br>(Alberta) | 192 | 52.9 | 74.6 | Not<br>reported | Hospital<br>clinic | Not reported | Not<br>reported | Not reported | Graduated High School 95%<br>Bachelor's Degree/higher 27.3% |
| Amtmann *et al.* (2014) | USA<br>(Greater<br>Washington) | 455 | 52.9<br>[10.8] | 82.9 | 91.4 | Community | RRMS 56.7%<br>Other 41.3%<br>Unknown 2% | Not<br>reported | Not reported | Did not graduate High School 0.4%<br>Graduated High School 13.4%<br>Technical School 8.8%<br>Some College 29.9%<br>College degree 29.5%<br>Advanced Degree 18% |
| Amtmann *et al.* (2015) | USA<br>(Seattle) | 164 | 53.2<br>[11.1] | 77 | 90 | Community | Not reported | Not<br>reported | Not reported | High School 15%<br>Some College/Associates Degree 49%<br>Bachelor's Degree 21%<br>Advanced Degree 15% |
| Patten *et al.* (2015) | Canada<br>(Calgary) | 152 | 50 | 77.6 | Not<br>reported | Hospital<br>clinic | RRMS 68%<br>SPMS 23.8%<br>PPMS 8.8% | Not<br>reported | No treatment 57.9%,<br>Glatiramer Acetate 25%,<br>Interferon Beta 11.8%<br>Fingolimod/Natalizumab 5.3% | Not reported |
| Gunzler *et al.* (2015) | USA (Ohio) | 3507 | 46 [12] | 73 | 83 | Hospital<br>clinic | RRMS 81%<br>CPMS 16%<br>Unknown 3% | Not<br>reported | Not reported | Not reported |
| Altura *et al.* (2016) | Canada<br>(Calgary) | 151 | 49.9<br>(24.9-82.3) | 76.8 | Not<br>reported | Hospital<br>clinic | Not reported | Not<br>reported | Not reported | High School/less 38.7%<br>Greater than High School 61.3% |
| Kim *et al.* (2017) | Unclear<br>(implied<br>USA) | 459 | 52.8<br>[10.9] | 82.6 | 91.5 | Community | RRMS 56.6%<br>Other 41.4%<br>Missing 2% | Not<br>reported | Not reported | Did not graduate High School 0.6%<br>Graduated High School 13.3%<br>Technical School 8.7%<br>Some College 30.1%<br>College degree 29.2%<br>Advanced Degree 18.1% |

## Findings on PHQ-9 performance indicators

### Appropriateness

Three studies(15,19,20) investigated the appropriateness of the PHQ-9 in pwMS with respect to its possible application for diagnosis, screening, and monitoring. Amtmann*et al. (2015)(15)* measured the appropriateness of using the PHQ-9 as a diagnostic tool for major depressive disorder (MDD) against criterion standard telephone administration of the Structured Clinical Interview for DSM-IV Disorders (SCID), concluding that it was inadequate for use in this role due to its Youden Index (YI) being substantially lower than 0.8 (observed YI = 0.55) even when the cut-off was optimised. In contrast, Patten *et al.* (19) used the same criterion method and reported data with an optimised YI of 0.83.

Patten *et al.* (19) primarily evaluated the appropriateness of the PHQ-9 as a screening tool for MDD, concluding that it performed well due to high sensitivity (95%) and specificity (88.3%) based on a cut-point of eleven. This was associated with a positive predictive value (PPV) of 51.4% and a negative predictive value (NPV) of 48.6%. Notably, use of only the first two items of the PHQ-9 (the PHQ-2) with a cut-point of three, provided sensitivity of 80% and specificity of 93%, with PPV of 64% and NPV of 36%.

Altura *et al.* (20) evaluated the appropriateness of the PHQ-9 as a screening tool for suicidal ideation, comparing the single PHQ-9 item on suicidal ideation against the criterion standard SCID item asking if the participant has had “recurrent thoughts of death, suicidal ideation, suicide attempt, or specific plan”. With a cut-point of one (any non-zero endorsement), the PHQ-9 exhibited sensitivity of 62.5%, specificity of 95%, PPV 41.7% and NPV of 97.8%. The high NPV was proposed as a basis to support suitability for use as a screening tool for the absence of suicidal ideation. No studies evaluated the appropriateness of the PHQ-9 for use as a monitoring tool for depressive symptomatology in pwMS.

### Reliability

Two studies investigated the reliability of the PHQ-9 in pwMS. Specifically, Amtmann et al. (2014)(14) evaluated the internal consistency of the PHQ-9 using item-total correlations, a technique where each item score is correlated with the summed score of all other items in the scale. An average item total score correlation was not reported, although the range was 0.35 to 0.67. The single PHQ-9 item below the commonly used ‘red flag’ threshold of 0.4 was that evaluating suicidal ideation. Unidimensionality of the PHQ-9 was nevertheless supported by 1-factor confirmatory factor analysis (n = 455) based on a comparative fit index (CFI) of 0.95 and Tucker-Lewis index (TLI) of 0.94. Sjonnesen *et al.* (18) reported similar item-total correlations ranging from r = 0.38 (suicidal ideation item) to r = 0.71, with an average of r = 0.55; Cronbach’s alpha was 0.82. No study evaluated the test-retest reliability of the PHQ-9 in pwMS.

### Validity

Four studies investigated the validity of the PHQ-9 in pwMS. Three explored the criterion validity of the PHQ-9 compared to the SCID criterion-standard in the context of their possible appropriateness to diagnose MDD(15,19), or to screen for MDD/suicidal-ideation(20). Separately, Amtmann *et al.* (2014)(14) evaluated the concurrent and discriminant validity of the PHQ-9. Large (concurrent validity) correlations were seen with the Center for Epidemiological Studies Depression Scale-10 (CESD-10; r = 0.85) and the 8-item PROMIS Depression Short Form (PROMIS-D-8; r = 0.73) depression scales. However, similarly high levels of correlation were also seen with the Modified Fatigue Impact Scale (MFIS; r = 0.73). Slightly lower (discriminant validity) correlations were reported between the PHQ-9 and both the PROMIS-Sleep disturbance (r = 0.57) and PROMIS-Pain interference scales (r = 0.60).

### Responsiveness

No study investigated the responsiveness of the PHQ-9 to clinically meaningful change in depressive symptomatology or quality of life in pwMS.

### Precision

Three studies (15,18,19) investigated the precision of the PHQ-9 in pwMS. The principal focus of this work has not been to remodel the fundamental structure of the PHQ-9 instrument (nine items with four response categories), but rather to explore the potential value of alternative scoring methods that aim to better reflect the relationship of item-specific responses to the underlying construct of depression. In particular, recognising the potential for some items to be differentially sensitive to ‘contamination’ by symptoms of MS *per se. Sjonnesen et al*. (18) hypothesised that PHQ-9 items evaluating fatigue and concentration deficits would be particularly prone to contamination, however no evidence was found to support exclusion of these items, or modification of the standard scoring system. Gunzler *et al.* (16) revisited this issue in a very large cohort (n = 3,507), using simultaneous measures of other MS symptoms in order to develop alternative weightings for PHQ-9 scores that maximised precision with respect to the construct of depression. The possible clinical utility of this adjusted scoring method has not yet been tested.

### Interpretability

The major focus of enquiry within this area has been on the meaning and value of established cut-points that were originally developed for generic use of the PHQ-9. Cut-points of 5,10,15, and 20 have been widely used to define ‘mild’, ‘moderate’, ‘moderately severe’, and ‘severe’ depression. This raises a fundamental question as to whether depression is best operationalised as a categorical or a quantitative state. The literature has primarily adopted a categorical approach. As context, large cohort (n = 580) evaluation in a general population of primary care patients identified a cut-point of 10 to have a sensitivity and specificity of 88% for MDD(16). In pwMS, Patten *et al.* (19) reported equivalent values of 95% and 85.9%, and Amtmann *et al.* (2015)(15) equivalent values of 93.8% and 61.2%. Using a group classification approach against SCID criterion standard, these two studies differed in their conclusion as to whether the PHQ-9 could be used as a diagnostic tool for MDD in pwMS. Amtmann *et al.* (2015)(15) reported a maximum YI of 0.55 even with optimisation of the cut-point (12), concluding that this was inadequate as it fell substantially below a minimum acceptable value of 0.8. In contrast Patten *et al.* (19) provided data indicating the YI did achieve this standard at a cut-point of 10 (YI = 0.809) and 11 (YI = 0.833), although not at a cut-point of 12 (YI = 0.756). Whether performance could be improved further at the individual clinical decision making level through alternative scoring systems such as those proposed by Gunzler *et al.* (16) has not yet been tested.

Kim *et al.* (17) investigated the potential for PHQ-9 scores to be interpreted with respect to their equivalent PROMIS-D depression scores, estimating these through a process termed ‘cross-walking’. The correlation between the direct PROMIS-D score and the ‘cross-walked’ PHQ-9 was moderately strong at 0.74. 56.6% of patients were categorized into the same PHQ-9 severity categories based both on actual and cross-walked scores, with 9.2% put into one lower category, 1.7% put in more than one category lower, 27.7% put one category higher and 4.8% classified into more than one category higher. Overall it was found that the PHQ-9 was most suitable for conversion to ‘cross-walked’ PROMIS-D scores in those with average to high depressive symptoms.

### Acceptability

No study explicitly investigated the acceptability of completing the PHQ-9 to pwMS. However, no substantial concerns were raised in prospectively recruiting studies where recruitment rate ranged from 28.8% to 98.1%, and retention/completion rate in longitudinal research was 80.9%.

### Feasibility

None of the studies explicitly investigated the feasibility of administering the PHQ-9 to pwMS. However, the literature includes a mixture of participants drawn from both primary and secondary care settings.

Taken together with the large total number of participants studied, this provides some evidence against there being substantial issues with feasibility in clinical practice.

## Discussion

### Summary of Evidence

We identified a modest literature of seven studies that specifically evaluated the psychometric properties of the PHQ-9 as a tool to measure depressive symptomatology in pwMS. The quality of these published studies was high and the overall number of participants reported on was substantial (n >5,000).

One-factor confirmatory factor analysis supported interpretation of the PHQ-9 as a unidimensional scale, consistent with measurement of a single underlying construct. Summation of PHQ-9 item scores is therefore a reasonable approach to provide a ‘global’ measure. Criterion validation of the summated PHQ-9 score against MDD, together with concurrent validation against other established ‘depression scales’ supports validity of the PHQ-9 as a measure for the underlying construct of depression. In contrast, discriminant validity findings have proved challenging to interpret. This difficulty reflects the fundamental lack of clinical features in MS that would be expected to vary independently, compounded by uncertainty about the extent and direction of any causal relationships between the severity of depressive symptomatology, fatigue, cognitive and physical impairments. Nevertheless, relatively high correlation with fatigue measures add to long-standing face validity concerns about the correct interpretation of PHQ-9 scores given the potential for overlap between the somatic features of depression and features of MS *per se*. This issue remains largely unresolved, and particularly problematic at the level of the individual for whom clinical decisions are required. It is possible that alternative scoring methods such as those proposed by Gunzler *et al.* (16) will provide more precise and clinically useful measurement, although this has not yet been fully evaluated.

The suitability of the PHQ-9 to be used as a diagnostic tool for MDD remains unclear, with the two studies that address this issue reaching opposing conclusions. In the absence of definitive evidence, it would therefore appear inappropriate to recommend use of only the PHQ-9 when making a diagnosis of MDD in pwMS. In contrast, value appears to exist for application as a screening tool for MDD. No consensus exists as to the optimum cut-off for use in this setting, however we favour a pragmatic approach based on maintenance of the widely used and readily recalled cut-off of ≥10 that provides a PPV of 51.4% and NPV of 48.6%, only marginally different from the optimised cut-off of ≥11 suggested by Patten *et al.*(19) Screening for suicidality based on the final item of the PHQ-9 is also effective, with a cut-off of ≥1 (*i.e*. any non-zero score).

Despite being a key property of any measurement instrument, scant data was available on the reliability of the PHQ-9. Acceptable internal consistency was demonstrated, however no information was available on test-retest reliability. Similarly, no data was reported on responsiveness to clinically meaningful change. Taken together, the suitability of the PHQ-9 for longitudinal use as a monitoring tool is therefore undetermined. Finally, the large number of participants and mixed setting of research environments can provide circumstantial support for the likely acceptability and its feasibility of use in clinical practice. However, it would be beneficial to formally evaluate these in subsequent research.

## Limitations

We have only been able to identify studies that have been published and so there may be a reporting bias. Greater uptake of study registration for observational research may mitigate this in future. Despite the large total sample size (n>5000), a single study provided 69% of the total cohort, and some concern was identified regarding the possibility that 39% of participants from the remaining studies may have been included in two reports. We nevertheless believe that the conclusions of these studies are not in question as they addressed distinct aspects of PHQ-9 performance. With regards generalisation of findings from the published literature, the female predominance (approximately 75%), overall mean age of approximately 50 years, and mixture of relapsing (c. 65%) and progressive MS disease types, raised no concerns about the representativeness of the overall cohort. However, we noted limited ethnic diversity, and that all studies were based on North American populations. Inclusion of greater ethnic and geographic diversity in future research would therefore be welcome.

## Conclusions

The PHQ-9 is a promising screening tool for MDD in pwMS and may have a role in diagnosis. However, significant gaps exist in the current evidence base around test-retest reliability, responsiveness, acceptability and feasibility that preclude conclusions regarding suitability for use as a depression monitoring tool in pwMS.

## Acknowledgements

We thank Siddharthan Chandran for helpful comments on the draft manuscript.

## Supporting information

**S1 Fig.** Systematic review protocol.

**S2 Fig.** Quality assessment tool for evaluation of manuscripts, based on the STROBE checklist.

